# Heterogeneous evolutionary history defines the rear edge of the North American herb *Campanula americana*

**DOI:** 10.1101/2025.11.24.690277

**Authors:** Antoine Perrier, Keric S. Lamb, Laura F. Galloway

## Abstract

**Aim:** Warmer range limits of species distributions offer a window into the ecological and evolutionary processes associated with long-term warming, as they often harbor relicts of refugial populations from last glaciation. However, warm-edge populations vary in their refugial, evolutionary, and climatic histories, complicating their use as models. Here, we characterize this variation at the warmer range limit of the North American herb *Campanula americana*.

**Location:** Eastern North America

**Taxon:** Campanula americana

**Methods:** We evaluated the warmer range limit using species distribution model hindcasting, range-wide analyses of population structure, effective migration and phylogenetic relationships.

**Results:** Warm-edge populations vary in their refugial history; most are relicts of historic refugial populations (i.e. rear edge), but some result from southward postglacial colonization. Within the rear edge, populations exhibit pronounced genetic structure, forming highly isolated clusters despite contiguous geography and only modest landscape barriers. Distinct rear-edge clusters also varied in climate history and gene flow among population, with the more southern populations experiencing higher habitat decline, fragmentation and isolation. Finally, only rear-edge populations from northern refugial areas contributed to postglacial range expansion.

**Main conclusions:** Our study highlights the challenges of using warm-edge populations as ecological and evolutionary models of response to climate change. Even in a geographically simple setting, warm-edge populations exhibit refugial, genetic, and climatic heterogeneity. Characterizing this variation is essential for accurately identifying rear-edge populations and inferring their evolutionary history. Knowledge of the contribution of warm-edge populations to range-wide genetic variation is key when considering how these populations may inform studies of adaptation to future climate change.

## Introduction

Understanding how changes in climate affect species distributions is a central challenge in ecology, evolution, biogeography, and conservation (Aguirre-Liguori et al., 2021; Lenoir & Svenning, 2015; Nadeau & Urban, 2019; Rubenstein et al., 2023; Urban, 2024). This challenge can be addressed by studying how historical processes have shaped range limits (Davis & Shaw, 2001; de Lafontaine et al., 2018). In temperate regions, Pleistocene glacial cycles repeatedly drove range contractions into refugia at low latitudes and elevations during cold periods, followed by expansions out of refugia in periods of warming (Hewitt, 2000; Hewitt, 2004). Much of the current range of temperate species thus reflects postglacial colonization (“expanded range”) after the Last Glacial Maximum (LGM, ∼20 kya; Hughes et al., 2013). Warmer range limits often harbor populations that have been exposed to near continuous warming since the LGM, and are thus increasingly being used to study ecological and evolutionary outcomes of warming (Perrier et al., under review in TREE; e.g. Masuda et al., 2023; Nielsen et al., 2024; Song et al., 2021; Tavares et al., 2024). However, these populations may vary in their refugial, evolutionary and climatic histories, both among and within species, and this complexity is underappreciated. Characterizing this variation is key for using warmer range limits as models of climate response.

Warmer range limits vary in whether they represent former refugial populations. In temperate species, postglacial range expansion generally occurred by tracking shifts in suitable habitats, expanding from warmer refugia into areas that were previously too cold (Hewitt, 2000; Hewitt, 2004; Stewart et al., 2009). Contemporary populations at warmer range limits may thus represent descendants of historic refugial populations that persisted more-or-less in place, and that have given rise to populations in the expanded range. These descendants are termed “rear-edge” populations (Hampe & Petit, 2005). The identity of rear-edge populations has often been assumed from their geographic position at the warmer latitudinal or elevational limit (Perrier et al., under review in TREE). However, warmer range limits do not necessarily conform to these expectations and may include expansion populations (e.g. Bouchard et al., 2017; Korábek et al., 2023; Willi et al., 2018). The assumption that warm-edge populations represent a rear edge, i.e. ancestral and occurring in refugia, must therefore be tested if they are to serve as models of response to past climate change. The location of refugia can be predicted by species distribution model (SDM) hindcasting, or from available fossil and pollen records, while ancestry can be tested with phylogeographic analyses (Gavin et al., 2014).

Rear edges are also not uniform, even within a single species. For example, temperate species may have multiple, geographically distinct rear edges, corresponding to separate refugia isolated by major geographic barriers (e.g. Aradhya et al., 2017; Assis et al., 2016; Hewitt, 2000; Provan & Maggs, 2011; Zeng et al., 2015). Separate refugia create opportunities for independent evolution and genetic divergence. Climate history may also vary among distinct rear edge regions. Refugia are often thought to have relatively stable climates (Gavin et al., 2014; e.g. Tzedakis et al., 2002), leading to long-term demographic stability with high genetic diversity in rear-edge populations (e.g. Erichsen et al., 2018; Neiva et al., 2020; Piotti et al., 2017). Alternatively, the continuous warming of refugial areas since the LGM may result in rear-edge populations occurring in areas where suitable habitats are rare and poor quality (Vilà-Cabrera et al., 2019), leading to genetic signatures of demographic decline and long-term isolation (e.g. Masuda et al., 2023; Sielezniew et al., 2015; Song et al., 2021; Tavares et al., 2024). Last, long-term environmental change at the rear edge may result in pronounced local adaptation (Bontrager et al., 2021), with unique adaptations to warming (e.g. Ghouil et al., 2020; Pelletier et al., 2023; Perrier et al., 2025). Understanding such diversity is important as it affects the extent to which inferences of vulnerability and adaptive potential of rear-edge populations are broadly applicable under future warming.

Finally, the genetic contribution of rear-edge populations to the expanded region of a species range may vary. Postglacial range expansion carries the assumption that refugial populations are ancestral to populations elsewhere in the range. By extension, the genetic diversity in expanded portions of the range is expected to be a subset of that in refugial populations. However, this may not always be the case. For example, expansion may have had a limited origin, such as from populations at the colder edge of refugia or from cryptic mid-range refugia (e.g. (Bemmels et al., 2019; Fernandez et al., 2021; Hantemirova et al., 2017; Lee-Yaw et al., 2008; Rull, 2009), with limited contribution from populations that now form the rear edge (Hampe & Petit, 2005). Alternatively, in species with multiple rear edges, some edges may contribute to expansion while others do not, with variation linked to the presence of geographical barriers to expansion. For example, in Europe high mountain ranges (e.g. Alps, Pyrenees) often divide refugia and expanded areas, limiting recolonization (Hewitt, 2004; Korkmaz et al., 2014; Petit et al., 2003). Evaluating the contribution of rear-edge populations to postglacial expansion is important for understanding the extent to which the rear-edge’s genetic diversity, and therefore evolutionary potential, may be shared with the rest of the range.

Characterizing populations at warmer range limits is important if we are to use them to understand response to past warming and to predict potential change to current climate warming. In this study, we explore the refugial, phylogeographic and climatic history of the warmer range limit of the North American herb *Campanula americana.* Specifically, we ask (1) Are populations at the warm edge of the range ancestral and located in refugial habitats, indicating rear-edge populations? (2) Does genetic complexity at the rear edge suggest multiple distinct refugia with potential independent evolution? (3) Have former refugial areas retained suitability or become marginal under past warming, and has this effected genetic patterns at the rear edge? (4) What is the contribution of rear-edge populations to postglacial range expansion? We find that rear-edge populations can be highly heterogeneous even in a fairly small and largely contiguous geographic area. Our study provides a framework for evaluating and interpreting the heterogeneity of warm-edge populations and for informing our understanding of how to use them as models of species responses to environmental change.

## Material and Methods

### Study system

*Campanula americana* is a monocarpic herb of open forests found in the eastern USA across a broad climatic and geographic range, from the subtropical Gulf coast to the Great Lakes, and from west of the Mississippi to the Appalachian Mountains (Fig. 1A). Populations belong to three geographically distinct genetic clades that diverged before the LGM, ca. 0.7 – 7.0 mya (Barnard-Kubow et al., 2015), and came into secondary contact along the Appalachian Mountains (Lamb et al., 2024). We focus on the larger “Western” clade, comprising most populations of the species (∼ 80%), and largely found at low elevations (<600m) west of the Appalachian Mountains. This clade is derived from the older “Appalachian” clade whose distribution is limited to the mountains. The Western clade is characterized by a history of rapid postglacial colonization with a mid-latitude origin estimated in eastern Kentucky (Barnard-Kubow et al., 2015; Koski et al., 2019; Prior et al., 2020). However, the phylogeography of the warmer range edge is unclear.

**Figure 1:**
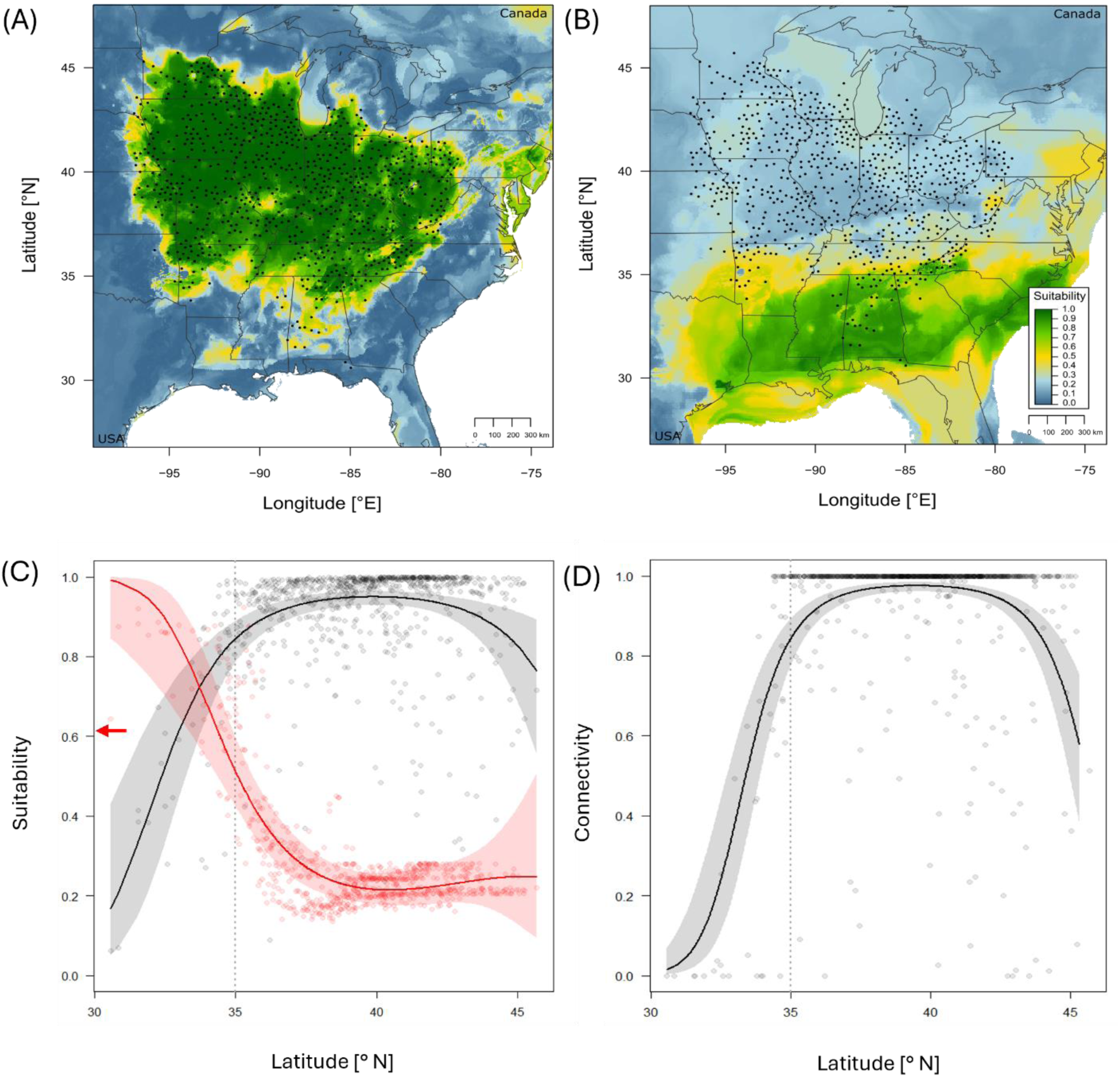
Climatic history of *Campanula americana*. Species distribution model (SDM) showing predicted habitat suitability during the present **(A)** and the Last Glacial Maximum **(B)**, with occurrences used to build the SDM as black dots. **(C)** Present-day (grey) and LGM (red) suitability, and **(D)** present day connectivity estimated for all occurrences used to build the SDM. Vertical dotted line represents the geographic separation between the rear edge (<35°N) and the expanded range. Lines represent the significant model-predicted relationship between each estimate and population latitude, with the 95% confidence interval indicated as shading. In **(C)**, the red arrow represents the present-day suitability threshold above which 95% of populations are included. Test statistics reported in Table 1.

**Table 1:**
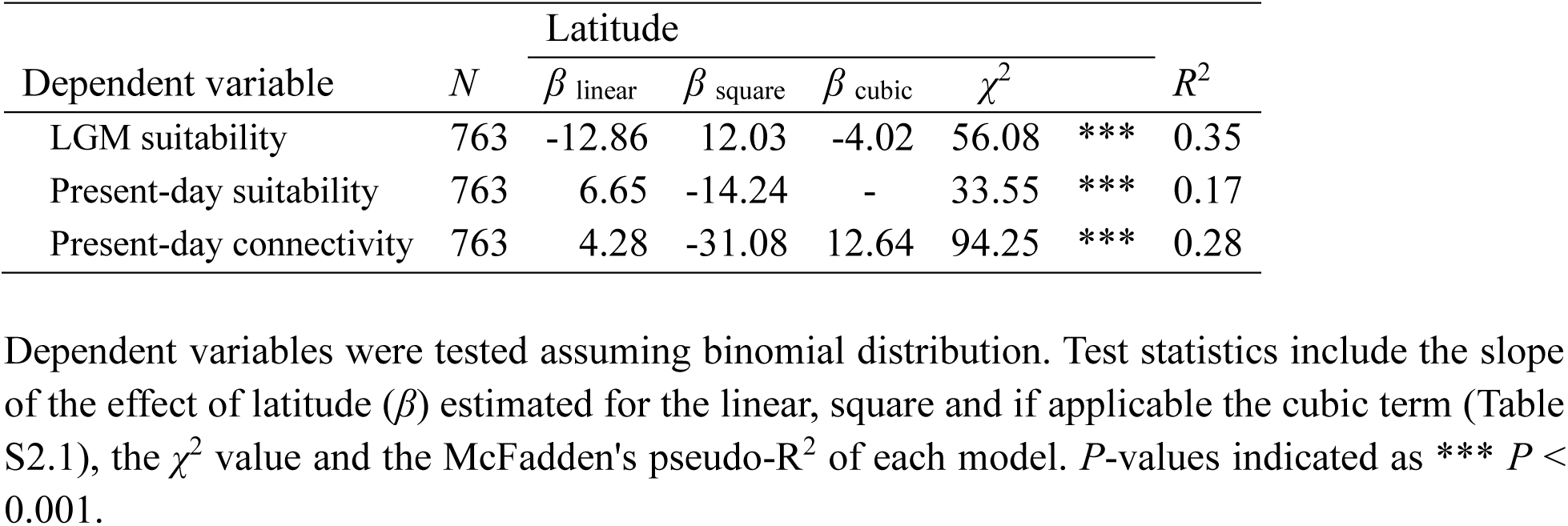
Test of variation in present-day and past habitat variables (Study 1) across the latitudinal range of *Campanula americana*.

We define the warmer edge of *C. americana*’s Western clade as roughly the lower latitudinal third of the range (below 35 °N, Perrier et al., 2025). This area is characterized by subtropical climates, with mild winters and rare freezing events, in contrast to the temperate climate found throughout the rest of the range. Warm-edge populations harbor unique phenotypes including populations that have lost the vernalization requirement for flowering (Perrier et al., 2025). Warm-edge populations broadly overlap with refugial areas and contain chloroplast haplotypes that, while not unique to this area, are ancestral to the clade (Barnard-Kubow et al., 2015), suggesting these populations are the rear edge of the species distribution.

### Species distribution model (SDM)

We created an SDM of populations in *C. americana*’s Western clade to identify potential glacial refugia though hindcasting (Q1) and to assess how these habitats have changed under postglacial warming (Q3). We first generated a SDM based on contemporary climates using Random Forest with the package *randomforest* (Liaw & Wiener, 2002) in R (R Core Team, 2024), based on 763 occurrences obtained from iNaturalist (Fig. 1A, www.inaturalist.org), and 19 bioclimatic variables obtained from WorldClim 2.1 representing a 30 year climate average (1970-2000; Fick & Hijmans, 2017). See Appendix S1 Method S1.1 in Supporting Information for details.

With this SDM, we developed a geographic projection of areas likely to have suitable climates based on the climate of present-day occurrences (WorldClim 2.1; Fick & Hijmans, 2017). Suitability is defined as the relative likelihood a species will be present at a location based on the climate, and ranges from 0 (unsuitable) to 1 (suitable). We defined suitable habitat for *C. americana* as areas with a suitability >=0.62, which corresponds to a suitability threshold that includes 95% of populations. To identify putative refugia (Q1), we assessed suitability across the range during the LGM by geographically projecting the SDM using downscaled paleo-climate data (i.e. hindcasting) as input (WorldClim 1.4, https://www.worldclim.org/data/v1.4/paleo1.4.html, accessed 04/11/2025). The location of glacial refugia was inferred as areas with suitable habitats during the LGM. Geographic projections of present-day and LGM suitability were then visually compared to assess differences in past and present suitability across the range (Q3).

We then estimated three habitat quality metrics for each iNaturalist occurrence from the SDM projections: LGM suitability, present-day suitability, and connectivity. Connectivity is a measure of habitat marginality and was estimated as the proportion of suitable habitat in a 25km radius circle around each iNaturalist occurrence, and ranges from 0 (no suitable habitats, i.e., low connectivity) to 1 (full area is suitable, i.e., high connectivity). We tested for differences in LGM suitability, present-day suitability, and present-day connectivity across the warm range edge and the expanded range in separate linear models, using latitude as a proxy for range position. Preliminary analysis tested whether latitude was best described by a linear, second-degree or third-degree quadratic relationship (the latter allows variables to vary at both high and low latitudes). Results are reported for the best model based on AICc scores (Table S2.1, Sugiura, 1978). If multiple models performed equally well (|ΔAICc| ≤ 2), the simplest model was chosen. Dependent variables were tested assuming binomial distribution. For each model (and in all subsequent analyses), we tested and confirmed model assumptions.

### Sequencing and genotyping

To determine phylogenetic relationships among populations (Q1, Q4) and population structure (Q2, Q3), we analyzed sequence data from 5 individuals in each of 36 populations (four for OK61, 179 individuals total, Fig. S2.1C, Table S2.2). These spanned the whole range of the Western clade of the species, excluding the Appalachian Mountains where gene-flow among clades may blur phylogeographic patterns (Lamb et al., 2024). We also included five individuals of Appalachian-clade population VA73 to serve as outgroup in phylogenetic analyses. Sequence data was generated using Restriction-site Associated DNA sequencing (RAD-Seq; Andrews et al., 2016; Baird et al., 2008) as part of a larger project described in Perrier et al., (under review in EVL; Bioproject PRJNA1306192). Details of the plant material, DNA extraction, library preparation, and sequencing are provided in Method S1.2. In short, DNA extraction was performed by the Genomics & Cell Characterization Core Facility at the University of Oregon (Eugene, OR, USA). Library preparation and sequencing were performed by Floragenex, INC (Beaverton, OR). Raw reads were processed into single nuclear polymorphism (SNP) calls following a modified STACKS v2 pipeline (Rochette et al., 2019) and using a haploid reference genome (Lopez-Caamal et al., under review in GBE). The species is an old autotetraploid that has undergone cytological diploidization (Lopez-Caamal et al., under review in GBE); prior phylogeographic studies have treated it as diploid (Barnard-Kubow et al., 2015; Koski et al., 2019; Lamb et al., 2024; Prior et al., 2020). Genotypes were thus called in diploid variant call format (VCF). SNPs were filtered for missingness (90%), minor allele count (5), sequencing depth (20x – 70x), and indels and linkage, resulting in a final set of 40,195 SNPs.

### Phylogeny and population structure

We assessed phylogenetic relationships among populations by constructing a phylogenetic tree with the Appalachian population VA73 as an outgroup using TreeMix v1.12 (Pickrell & Pritchard, 2012). TreeMix was chosen over other population-level phylogenetic methods to account for the admixture identified in the population structure analysis (see below). TreeMix was run assuming one to 10 migration events (m). For the best (m), TreeMix was run again with 100 bootstraps and visualized following the workflow of the *BITE* package in R (Milanesi et al., 2017).

We inferred the genetic structure across 36 populations by determining individual-level genotype clustering using sparse nonnegative matrix factorization (sNMF) implemented in the *LEA* package in *R* (Frichot & François, 2015). The sNMF was run assuming one to 36 clusters (*K,* details in Method S1.2). Individual-level admixture, the proportion of an individual’s genome assigned to genetic clusters, was generally consistent within a population (Fig. S2.1B). We therefore focused on population-average admixture and assigned a main genetic cluster to each population (Method S1.2, Table S2.2).

We characterized geographic patterns of gene flow to complement the population structure analysis by estimating effective migration surfaces (Petkova *et al*. 2016) implemented in the python package *feems* (Marcus et al., 2021); Method S1.2). This method identifies geographic areas where gene flow between populations is higher or lower than expected under isolation-by-distance. It provides a spatially explicit visualization of population structure, allowing for identification of geographic barriers and corridors of gene flow.

To infer the colonization history of the species, we visually compared the results of the three phylogeographic analyses with the distribution of past and present-day suitability. We also compared these results with large geographical features across the range that could serve as barriers to infer their potential impact on colonization history and population structure.

## Results

### Identification of glacial refugia

The SDM was a good match for the current range of the species (Fig. 1A), with a small error rate of 2.1% false negatives and 4.7% false positives, indicating it captures the climatic niche of the species well. Therefore, the SDM is appropriate for hindcasting the niche of the species to the LGM. Hindcasting the SDM revealed that the contemporary range above ∼ 35 °N was unsuitable during the LGM (Fig. 1B), indicating that populations in this area likely established after the LGM by tracking shifts in suitable habitats. In contrast, the lower latitudinal range, below ∼ 35 °N, overlaps with an area of high LGM suitability extending from the northern Gulf coast up to the southern Appalachians, indicating it may have served as refugia during the LGM. Therefore, contemporary populations at *C. americana*’s warmer range edge occupy likely refugial habitats and thus may be remnants of refugial populations that have persisted more-or-less in place until the present day. LGM habitat suitability showed a significant third-degree quadratic relationship with latitude (Table 1, Table S2.1). Suitability was greatest at low latitudes and declined rapidly towards higher latitude, with LGM habitats becoming unsuitable (<0.62 suitability) above ∼34.5 °N (Figure 1C), supporting inference from geographic projections.

### History of postglacial climate change

Warmer range edge habitats have deteriorated since the LGM. The SDM projection indicates that most populations across the species’ distribution currently occur in suitable habitats, while southern parts of the range are less suitable and become increasingly fragmented (Fig. 1A). In line, present-day habitat suitability and connectivity both had significant negative quadratic relationships with latitude (second-degree and third-degree, respectively, Table 1, Table S2.1), with the highest values at mid latitudes and sharp declines towards low latitudes, particularly below 35 °N (Fig. 1C, D), and to a lesser extent at high latitudes. This relationship predicts that contemporary habitats become unsuitable (<0.62 suitability) below ∼33°N. Only a small area along the northern edge of the putative refugia retain suitability today (Fig 1A, B).

### Phylogeography of the rear edges of C. americana

Most warm-edge and refugial populations belong to a distinct early-diverging clade. For the best migration (m=1) and topology, the phylogenetic tree largely recapitulated geography and shifts in habitats between the LGM and present-day. Populations belong to two major and geographically distinct clades (Table S2.2, Fig. 2A). The “Refugia” clade is largely composed of populations occurring in potential refugia identified in the SDM analysis, except for the most basal one, TN2, occurring just beyond the northern edge of the refugia. In contrast, most populations of the Expansion clade occur in areas unsuitable during the LGM. The exception, OK61, occurs in the refugial area just south of the Ozark mountains, and is one of the most derived expansion populations, suggesting relatively recent recolonization of refugial habitats. The overall topology, and the split between the Refugia and Expansion clades, was well supported by bootstraps. Within each clade, bootstrap support varied and was particularly low for the basal branches of the Refugia clade.

**Figure 2:**
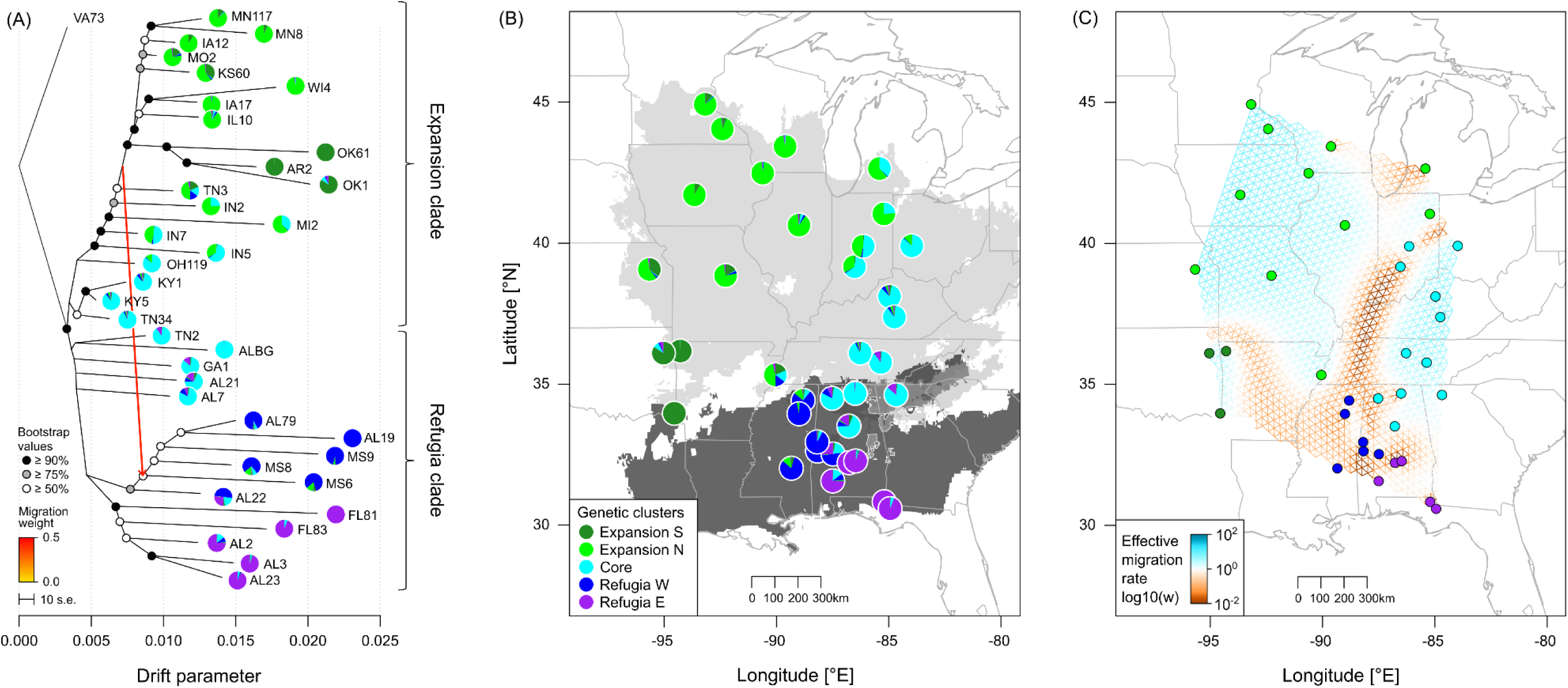
Phylogeography of *Campanula americana*. **(A)** Phylogeny based on nuclear data (VA73 as outgroup) with pie chart colors indicating genetic clusters and polymorphism within populations. Brackets detail which populations belong to the Expansion and Refugia clade. Migration event and its inferred direction are displayed as a colored arrow (see scale). The bootstrap value of each branch is indicated for values ≥ 50%, estimated from 100 bootstraps. The scale of 10x the standard error for the drift parameter is indicated. **(B)** Sequenced populations with genetic clustering as in (A). Light grey shading represents contemporary suitable habitat, dark grey represents the location of putative refugia, with intermediate hues representing areas suitable both during LGM and today. **(C)** Estimation of the effective migration surface between populations with populations colored by main genetic cluster. Areas where gene flow is lower than expected under isolation-by-distance are represented in brown, areas where gene flow is higher in blue.

Warm-edge populations have substantial genetic structure. Range-wide genetic variation was best partitioned into five genetic clusters in the population structure analysis (best K = 5, best run = 3, Method S1.2, Fig. S2.1A). The genetic clusters are geographically structured (Fig. 2B, Table S2.2). Refugial populations in the southeastern part of the range belong to two “Refugia” clusters separated longitudinally (Refugia W, Refugia E). Further north, populations belong to a “Core” cluster that spans the putative refugial habitat and the expansion region of the range. Populations in the expanded western part of the range belong to two clusters, the large Expansion N cluster in the northwest and the smaller Expansion S in the southwest of the range. Patterns of population structure generally recapitulated the phylogeny (Fig 2A). The “Refugia” clade is largely composed of populations from the two Refugia clusters, while the “Expansion” clade is mainly composed of populations from both Expansion clusters. Populations from the Core cluster are basal in both clades with the five southernmost populations at the base of the Refugia clade, and the six northernmost populations at the base of the Expansion clade. Both clades show substructure reflecting genetic clusters, with populations from the Expansion S and from the Refugia E and W forming distinct sub-clades.

Most refugial populations show high isolation among and within genetic clusters. The effective migration analysis identified limited migration between the clusters characterized by the population structure analysis (Figs. 2B, C), with all clusters separated by areas of low gene flow. Within the Expansion clusters, populations were connected by areas of high gene flow, except for the northeast most population (MI2) in Expansion N. Core populations were also generally well connected, except the three southernmost. In contrast, the two Refugia clusters showed reduced gene flow between populations within clusters.

Admixture between genetic clusters was common and largely reflects geographic proximity and ancestry (Fig. 2B). Populations across the range showed large variation in admixture between genetic clusters, with the cumulative proportion of genetic variation assigned to secondary clusters ranging from 0.00 to 0.54 (mean = 0.17, Method S1.2), Admixture was homogeneous at the individual level (Fig. S2.2B), suggesting potential gene exchange is old and not ongoing. Highly admixed populations mostly formed between geographically and phylogenetically close clusters and occurred at the geographic boundaries between clusters (Fig. 2C). These patterns were also found in the effective migration analysis, evidenced by small breaks in the areas of low gene flow between clusters (Fig. 2C). One notable exception was admixture between southern populations of the Expansion N (TN3) and S clusters (OK1) and three populations of the Refugia W cluster (MS6, MS8, MS9; Fig. S2.2B). This was also identified in the phylogeny as a large admixture event from the Expansion clade into the Refugia clade (Fig. 2A).

### Link between phylogeographic patterns and landscape features

Phylogeographic patterns partially coincide with the extent of the LGM ice sheets. The three most basal populations of the Expansion clade (KY1, KY5, TN34; Fig. 2C) are located just south of the ice sheet (Fig. 3) and form a distinct sub clade. In contrast, the next most derived populations along the phylogeny occur just to the north in areas covered by ice sheets during LGM (Fig. 3), suggesting range expansion after the retreat of the ice sheets.

**Figure 3:**
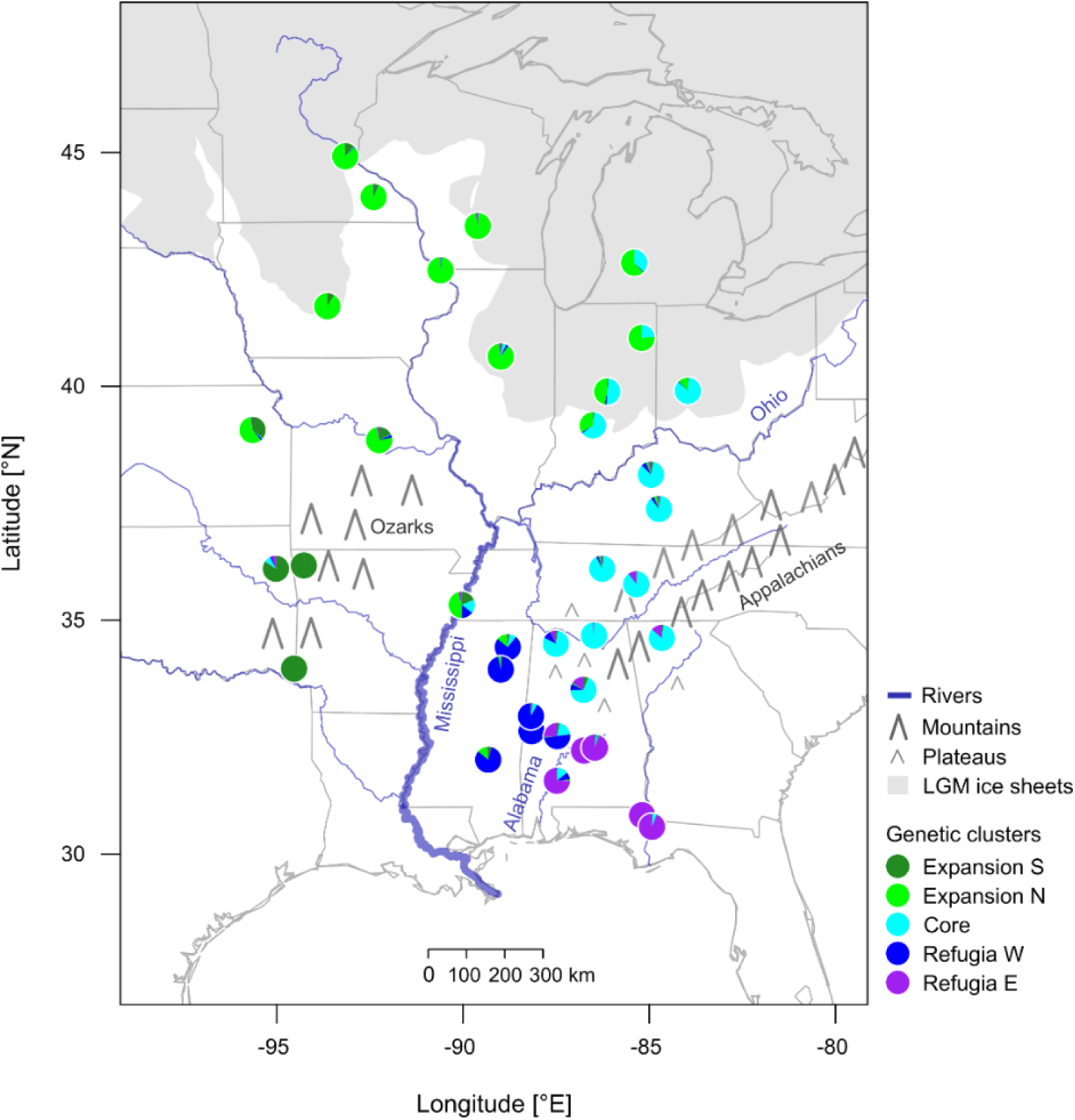
Geographic features across the range of *C. americana*. Location of the sequenced populations (see Fig 2A), with the extent of the ice sheets during the Last Glacial Maximum (gray shading), major rivers crossing the sampled range (blue), and relevant mountains (gray triangles).

Landscape features align with phylogeographic boundaries in the expanded range, but less so at the warmer range edge (Fig. 3). The Ohio and the lower Mississippi rivers, separate both Expansion clusters from the rest of the range. The Expansion S cluster is partially separated from the Expansion N cluster by the Ozark Mountains. The Refugia clusters are separated by the Alabama river. In contrast, the Refugia and the Core clusters are not separated by defining geographic features, though they differ in physiographic regions. The southern Core populations occur in mountainous areas including the southern reaches of the Appalachian Mountains and uplands such as the Cumberland plateau and the southern Appalachian Piedmont, while Refugia population clusters occur in the Coastal plains.

## Discussion

### Refugial clades and southward expansion lead to a complex warmer range limit

The genetic composition of species’ warmer range limits may be heterogeneous, including populations that vary in their refugial, phylogeographic, and climatic history. Most are expected to be rear-edge populations, relicts that persist in glacial refugia and are ancestral to populations derived from postglacial expansion. However, warmer edges and rear-edges of species distributions are often conflated (Perrier et al., under review in TREE), with “rear edges” identified by geographic position (i.e. equatorial edge), climate (i.e. warm edge), refugial overlap or phylogeographic history (i.e. basal). Depending on which criterion, or combination of them, is applied, different populations may be designated as the rear edge. We found that most populations at *C. americana*’s warmer range limit represent persistent refugial populations, i.e. stable rear edges (*sensu* Hampe & Petit, 2005). Hindcasting SDM revealed that contemporary populations within the geographic rear edge (<35 °N) largely occur in putative LGM refugia along the northern Gulf Coast. These refugia are consistent with earlier findings in the species (Barnard-Kubow et al., 2015) and with proposed refugia for other Eastern North American (ENA) plants and animals (Lyman & Edwards, 2022; Soltis et al., 2006), particularly species with distributions similar to *C. americana* (Bemmels & Dick, 2018; Mohn et al., 2021; Walker et al., 2009; Wang et al., 2018; Wyatt et al., 2021; Xia et al., 2024). Populations occurring in refugial areas and east of the Mississippi also form a distinct Refugia clade that diverged early from populations restricted to areas unsuitable during the LGM. The combination of their deep phylogenetic divergence and persistence in long-term refugia supports the inference that these warm-edge populations represent *C. americana*’s rear edge.

Postglacial recolonization can lead to heterogeneity in warmer range limits, even among populations in refugial areas. For example, the southwestern OK61, although located in a putative refugial area, derives from northern populations of the Expansion clade, suggesting post-LGM recolonization of refugia. Therefore, this population does not represent the rear edge. Unlike the Refugia clusters, it inhabits a relatively small putative refugial area west of the Mississippi. While postglacial expansion into warmer areas may seem counterintuitive, similar cases have been reported across the globe, including plants in ENA (Bouchard et al., 2017; Willi et al., 2018) and land snails in Europe (Korábek et al., 2023). We identified additional genetic signatures of the Expansion clade in refugial areas through an admixture event from the southern reaches of *C. americana*’s Expansion clade into the western-most populations of the Refugia clade. Together, these findings indicate that populations in warmer range limits are not always ancestral, even if they occur in putative refugia, underscoring the importance of testing the assumption that warm edges are synonymous with rear edges.

### Multiple contiguous but isolated refugia lead to high rear-edge population structure

The rear edge of *C. americana* is highly structured despite occurring in a relatively restricted geographic area with limited topographic heterogeneity. It is composed of three largely independent genetic clusters – the southern populations of the Core cluster and the Eastern and Western Refugia clusters. These clusters are highly differentiated and isolated from one another, despite geographic proximity, suggesting that populations were confined to three independent refugia with limited gene flow among them during the LGM. In contrast, most of the expanded region of the Western clade has little structure and is occupied by a single cluster (Northern Expansion) with the Southern Expansion cluster modest in size.

Modest geographic features appear to have led to isolation between the distinct yet neighboring rear-edge clusters. The Refugia clusters are separated by the Alabama River. Populations along this border show some admixture suggesting only partial isolation from the river. Rivers in this region experienced substantial changes during the late Pleistocene (Leigh, 2006), likely affecting their capacity to act as barrier. Local topography may have separated the southern Core cluster populations from the refugia clusters. Habitats at the southern extent of the Core cluster consist of plateaus separated by deep valleys (e.g. Cumberland Plateau, Ridge and Valley, Appalachian foothills). Limited gene flow among the southernmost populations of the Core cluster supports these physical features as barriers contributing to local isolation. In contrast to these modest geographic barriers, prominent present and past geographic features contribute to the delineation of genetic clusters in the expanded range, such as the Ohio River and the southern extent of the LGM ice sheet (Core-Northern Expansion break), and the Ozark Mountains (Northern-Southern Expansion break).

Multiple genetically distinct and isolated rear edges are not uncommon. However, they typically involve well-separated refugia, divided by large distances or prominent landscape features. European species with refugia isolated in distinct Mediterranean peninsulas are classic examples (Havrdová et al., 2015; Jiménez-Mejías et al., 2012; Korkmaz et al., 2014; Petit et al., 2003; Svenning et al., 2008). In ENA, major landscape features such as the Mississippi river and the Appalachian Mountains may also isolate refugia, but have typically been associated with breaks between distinct clades that diverged well before the LGM (Lyman & Edwards 2022; Soltis et al. 2006). In contrast, our study found the development of strong rear-edge genetic structure within a single clade and in a restricted geographic area, even in the absence of large-scale geographic barriers. This pattern is unusual, especially at such a small geographic scale and suggests greater attention is needed to the phylogeography of rear edges.

### Rear-edge populations are isolated

Populations in the rear edge are more isolated from each other than populations elsewhere in the range. At the rear edge, gene flow is limited among populations within clusters relative to expectations under isolation by distance. In contrast, populations in the Expansion clade, including northern Core populations and the Expansion clusters, have high levels of gene flow within clusters. The difference is particularly striking given that rear-edge populations were sampled more densely than the rest of the range. In the Expansion clade, substantial gene exchange within clusters but limited gene exchange between clusters that arise sequentially along the phylogeny suggests a serial range expansion process, with infrequent crossings of major geographic barriers and largely unobstructed gene flow within regions. This history contrasts with the rear edge, where populations did not experience range expansion and there are few barriers to explain isolation among populations within clusters. This pattern suggests other mechanisms underlie the high isolation at the rear edge.

Sustained postglacial warming over long periods of time may have led to isolation among rear-edge populations. While rear-edge habitats were once almost uniformly suitable for *C. americana*, present-day habitats show a steep reduction in suitability, especially towards the southern range limit where patches of suitable habitat are scarce and have low connectivity. This is expected to lead to small, isolated populations, expectations supported by the limited gene flow among populations in the southern clusters relative to expectations of isolation by distance. Therefore, progressive habitat decline and fragmentation under postglacial warming may have driven long-term isolation among rear-edge populations despite their geographic proximity. This pattern may be common in temperate taxa where rear-edge populations frequently occur in ecologically marginal habitats (Perrier et al., under review in TREE; Vilà-Cabrera et al. 2019), near species’ thermal and xeric physiological limits (Cahill et al., 2014).

Not all rear-edge habitats have been equally effected by postglacial warming. The northern part of the refugium has higher contemporary suitability and connectivity than the southern portion, indicating more modest postglacial warming with more habitat remaining within *C. americana*’s environmental tolerance. In line, northern refugial populations also show somewhat less isolation than more southern rear-edge populations where suitability and connectivity are low. Variation in evolutionary response to past warming has been documented across several taxa, but often only among geographically distinct rear edges (e.g. Havrdová et al., 2015; Neiva et al., 2014; Provan & Maggs, 2011). We find within a single, contiguous refugial region, rear-edge populations can differ in their climatic histories, with lasting consequences for patterns of genetic variation among populations

### The contribution of the rear edge to range expansion

A subset of rear-edge populations contributed to postglacial range expansion in *C. americana*. Only the southern populations in the Core cluster share ancestry with the expanded range and therefore served as the source of range expansion. The Refugia clusters did not contribute to range expansion. While variation in the contribution to range expansion among rear-edge populations is common (Hampe & Petit, 2005), it is typically found among geographically distinct rear edges (Hewitt, 2000; Lee-Yaw et al., 2008; Petit et al., 2003; Zeng et al., 2015), where the presence or absence of geographical barriers limiting poleward range expansion is associated with ancestry of populations in the expanded range (Korkmaz et al., 2014; Petit et al., 2003). Unglaciated ENA lacks comparable barriers, and most geographical features that have been associated with phylogenetic breaks run north to south (Lyman & Edwards, 2022; Soltis et al., 2006). This suggests a relatively permissive corridor for range expansion (Bemmels & Dick, 2018; Park & Donoghue, 2019), with uniform expected contribution throughout the rear edge. Despite this, we find that rear edges may be composed of populations that differ in their contribution to genetic variation in the expanded range, and that this variation may be observed even among neighboring populations.

The origin of postglacial expansion remains unclear in *C. americana*. Populations in northern, formerly unsuitable habitats, form a monophyletic clade, with northern and western populations at its tips and central populations in the east at its base (Tennessee, Kentucky), consistent with rapid westward range expansion from a mid-latitude origin (Koski et al., 2019; Prior et al., 2020). However, the colonization origin is north of the presumed refugia. One explanation for an origin in apparently unsuitable habitat is possible northern microrefugia, not identified by the SDM (see discussion in Prior et al., 2020). Postglacial colonization from microrefugia in colder areas than main refugial habitats is common among temperate taxa (Gavin *et al*. 2014; Rull 2009; e.g. Fernandez et al., 2021; Hantemirova et al., 2017), including ENA species (Peterson & Graves, 2016). Alternatively, the shared ancestry between northern Refugia-clade populations and Core-cluster populations that are basal to the Expansion clade implicates northern parts of the refugia as the origin of range expansion. In this case, the mid-latitude origin of the Western clade would represent a secondary expansion. In species with multiple refugia, greater contribution to postglacial colonization from colder refugia has also often been reported (e.g. Lee-Yaw et al., 2008). Our findings highlight that the pattern of colonization from colder areas also holds in the case of geographically contiguous refugia.

Which Western-lineage *C. americana* populations are oldest? In species or lineages that experienced postglacial range expansion, the oldest populations should be found at the rear edge (Hampe & Petit, 2005; Park & Donoghue, 2019). However, in contrast to the expected south-to-north gradient of ancestry (Barnard-Kubow et al., 2015), the base of *C. americana*’s Western clade is nested within the Core cluster in the phylogeny. The Core cluster populations are broadly located in the southern Appalachian Mountains where the Western clade of the species is thought to have emerged during the mid-Pleistocene (0.7-2.3 mya; Barnard-Kubow et al., 2015). One possibility is that the Core cluster is the closest to early Western populations, and that southern rear-edge populations represent an early split from this ancestor during (or before) LGM. The poor bootstrap support of southern core populations suggest that incomplete lineage sorting may obscure a clear phylogenetic signal of this history. This is especially likely if populations in the core remained large and connected (Maddison & Knowles, 2006). Indeed, we found high gene flow among populations in the core of the range, where refugial populations occur in climates that are also suitable today, suggesting demographic stability through time. Overall, these phylogenetic patterns highlight that rear edges differ not only in their contribution to range expansion but also in their ancestry, and suggest that distinct demographic and climatic histories contribute to their phylogenetic complexity.

### Integrating rear edge complexity in models of climate response

Characterizing the multiple dimensions of variation in rear-edge populations is key for using them as ecological and evolutionary models. We found that not all warm-edge populations represent rear edges. Distinguishing evolutionary history is important because populations in refugia that result from recent range expansion may have reduced diversity due to serial founder events and bottlenecks relative populations that persisted in the rear edge since the LGM (Excoffier et al., 2009). Similarly, populations with a long history of postglacial warming may have evolved adaptations to the warm climate while more recent arrivals may be less well adapted or have a different set of adaptations. Further, gene flow from populations in the expanded range into the rear edge may reduce fitness, swamping adaptation to warm climates, or, alternatively, increase fitness by providing novel variants (Aguirre-Liguori et al., 2021; Kottler et al., 2021; Sexton et al., 2011). Finally, the substantial genetic structure of the rear edge, with genetically distinct populations and clusters, provided the opportunity for independent evolution in response to postglacial warming, yielding possible distinctive adaptive outcomes (e.g. Havrdová et al., 2015; Neiva et al., 2014; Provan & Maggs, 2011). With this breadth of variation, assessing the diversity of the rear edge is key for using them as models of response to warming climates and to infer their potential contribution to ongoing change.

Finally, we found that not all rear-edge populations contribute equally to postglacial range expansion, even among neighboring populations. Rear-edge populations often exhibit strong local adaptation (Bontrager et al., 2021), with unique traits that enable persistence under warming climates (e.g. Ghouil et al., 2020; Pelletier et al., 2023; Perrier et al., 2025). If the underlying genetic variation is shared across the range, these adaptations could facilitate response to future warming in northern populations. However, if rear-edge populations did not contribute to range expansion, their adaptive variation may be regionally confined and contribute little to range-wide responses. For example, in *C. americana*, warmer winters reduce reproductive success in northern populations but have little effect on fitness in southern rear-edge populations (Perrier et al. 2025). The limited contribution of these southern rear-edge populations to the expanded range raises concerns about the capacity of more northern populations to adapt to future warming. Identifying which populations historically contributed to range expansion, and which did not, will be essential for using rear-edge populations as models of range-wide responses to climate change.

## Acknowledgements

This work was supported by the Swiss National Foundation (P2BSP3_195363), the National Science Foundation (NSF DEB-2140189), and the University of Virginia College of Arts and Sciences. For their help in raising of plants, we thank C. Claussen, S. Cox, O. Keenan, A. López, H. Makowski, and M. Turner. We are also grateful to D. Brown, J. Collins, A. Diamond, L. Elliott, K. England, E. Galloway, F. Griffith, I. Guenther, J. Hansen, B. Hoagland, H. Horne, J. Kees, M. Kohout, R. Laporte, L. Michael, D. Reed and B. Sutherland for seed and leaf material collection in natural populations. Collection permits were provided by the Florida Department of Environmental Protection, the Tennessee Department of Environment and Conservation, and Missouri Department of Conservation.

## Conflict of Interest Statement

The authors declare that there is no conflict of interest.

## Author Contributions

All authors contributed to the study design. KL built the SDM, and AP analyzed the SDM output. KL and AP collected seeds in the field and raised plants for sequencing. AP prepared samples for sequencing, processed and analyzed the sequencing data. AP wrote the manuscript with input from LG and KL.

## Data availability statement

Following data is available on Zenodo (https://doi.org/10.5281/zenodo.17704032): Presence-absence data to generate the SDM, raster layers of Present and LGM projections of the SDM, suitability and connectivity data derived from these projections, list of sequenced samples, individual genetic cluster assignments, the code to filter the VCF and the final VCF. Read data are publicly available in the Sequence Data Archive (BioProject PRJNA1306192).

## Notes

### Competing Interest Statement

The authors have declared no competing interest.

