## Appendix 1 for "Heterogeneous evolutionary history defines the rear edge of the North American herb *Campanula americana*"

**Appendix 1: Supplementary Methods**

**Method S1.1: Species distribution model**

*Data curation and pseudo-absence generation*

To model the current and historic range of the western cluster of *C. americana,* we first downloaded all research-grade observations for *C. americana* available on iNaturalist (7780 records, www.inaturalist.org, accessed 01/05/2022). To reduce sampling bias in where naturalists record the species, we thinned observations to a minimum spacing of 25km. We also excluded observations with fewer than 5 neighbors within 100km to remove isolated observations far beyond the known range limit.

Observations were then pruned to represent the distribution of *C. americana*'s Western clade. We assembled a set of records of populations known to belong to this clade based on previous studies in the species (Barnard-Kubow et al., 2015; Koski et al., 2019; Lamb et al., 2024), and generated a concave hull with a 100km buffer around these populations. We then pruned iNaturalist observations keeping those within the hull, to create a final set of 763 observations (Method S1.1 Figure).

Next, we generated a set of randomly located pseudo-absences. We first created a dense (0.2 °N x 0.2 °E) and randomly jittered (± 0.1°) grid of latitude (30 to 50 °N) and longitude (-100 to -70 °E) coordinates extending beyond the range to serve as pseudo-absences. We then restricted these pseudo-absences to those that fell within 200km of any given observation. Next, we generated a hull around all observations with a 50km buffer and removed any pseudo-absence points generated within this buffered hull. This filtering was performed to reduce the risk that pseudo-absences are generated in habitats where *C. americana* is present, but naturalists have not recorded it, or that pseudo-absences are generated in habitats within *C. americana*'s fundamental niche but it is not observed. Finally, we thinned pseudo-absences to a minimum spacing of 25km, then randomly selected an equal number of pseudo-absences as observations to create a dataset to use in modeling (Method S1.1 Figure).

*Modeling*

We used a Species Distribution Model (SDM) to estimate present-day suitability of sites for *C. americana*. We initially extracted the 19 bioclimatic variables from the WorldClim 2.1 1970-2000 climate data (2.5 arc-minutes resolution; Fick & Hijmans, 2017) for all observations and pseudo-absences. We then performed a PCA on this bioclimate data to reduce dimensionality accounted for correlation among bioclimate variables. We then used the first five principal components (> 95% of cumulative variance) as predictor variables in the SDM. We built a SDM with Random Forest implemented in the *randomforest* R package (Liaw & Wiener, 2002). To test how the model performed, we first trained a model on a dataset comprising a randomly sampled

70% of the data. We then tested how often the model predicted an absence where an observation existed in the remaining 30% of the data (2.1% false negatives), and how often the model predicted a presence where a pseudo-absence had been generated (4.7% false positives). The final SDM was then generated using all data, with 5,000 trees and a node size of 10.

*Predicting past and present distributions*

To model current distributions, we predicted present-day range-wide patterns of habitat suitability. We generated a dense (0.05 °N x 0.05 °E) grid of latitude (25 to 48 °N) and longitude (-100.5 to -66.5 °E) coordinates, then extracted all WorldClim 2.1 bioclimate data for these coordinates. This data was converted to PC coordinates based on the PCA generated with the observations and pseudo-absences. We then predicted range-wide patterns of habitat suitability from the SDM using the function predictSDM from the *mecofun* R package (Zurell, 2020). This data was then converted to a raster (WGS84) for visualization and downstream analysis.

To model past distributions, we hindcasted the SDM onto paleoclimates to predict range-wide patterns of habitat suitability during the Last Glacial Maximum. We extracted paleoclimate data (~22kya) from the WorldClim 1.4 Global Climate Model (CCSM4 model, 2.5 arcminutes resolution; https://www.worldclim.org/data/v1.4/paleo1.4.html, accessed 04/11/2025) for the same grid as generated for the present-day climate. The paleoclimate data was converted to PC coordinates as described above and used to predict LGM habitat suitability.

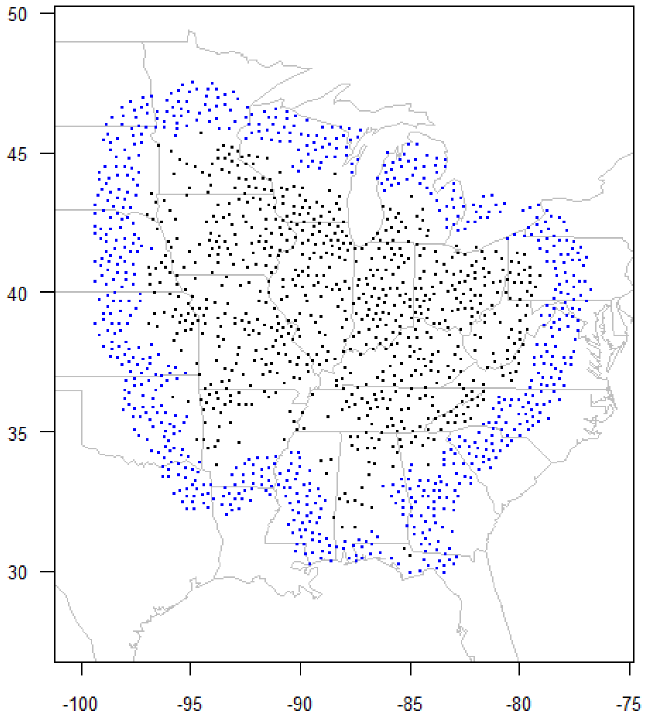


**Figure S1.1:** Final set of presence (black) and pseudo-absences (blue) used to generate the species distribution model.

**Method S1.2: Phylogeography**

We analyzed nuclear sequence data from 184 individuals representing 36 populations (Table S2.2) across *C. americana*'s Western clade, and one population (VA73) from its Appalachian clade to serve as outgroup (five individuals per population, four for OK61). These samples were part of a larger sequencing project described in Perrier et al., (under review in EVL, Bioproject PRJNA1306192). In that project, up to 10 individuals were sequenced for 31 of the populations (included in Perrier et al., under review in EVL). To ensure equal representation across populations in the present study, we retained only the five individuals with the highest coverage from those 31 populations. This sampling also reduces missing data and allows consistent comparisons in phylogenetic and population structure analyses.

*Plant material*

Plant material was sampled from natural populations or greenhouse grown plants. Most leaf material was collected from plants grown from seeds following one generation of random within population-crosses in the greenhouse (111 individuals, Perrier et al., under review in EVL), with the remainder from greenhouse-raised seeds from presumably unrelated plants in natural populations (63 individuals) or leaf material sampled from nature (10 individuals). Samples were dried in at low heat (~35 to 40 °C) for two to three days, then preserved at -20 °C.

*DNA extraction, library preparation & sequencing*

DNA extraction was performed by the Genomics & Cell Characterization Core Facility at the University of Oregon (Eugene, OR, USA), and RAD-Seq library preparation and sequencing were performed by Floragenex, INC (Beaverton, OR). For each sample, DNA was extracted from ~ 20mg of dry leaf tissue (MagMAX™ Plant DNA Isolation Kit, Applied Biosystems, Foster City, CA, USA) then purified (Mag-Bind® TotalPure NGS, Omega Bio-tek, Norcross, GA, USA). Individual samples were then concentrated, tested for purity and fragmentation, and equilibrated to 20 ng/µL. Samples were processed in five batches of 95 individuals from extraction to sequencing (475 total in the larger project). Individual DNA samples were digested by the *PstI* restriction endonuclease, and resulting fragments were tagged with multiplex RAD adaptors (95 unique 10 bp barcodes). Fragments were pooled by groups of 95 (multiplexed), sheared by sonication, size selected (300–500 bp range) and amplified by PCR. Each library was paired-end sequenced (150bp) on two NovaSeq6000 S4 flow cell lanes (Illumina, Inc. San Diego, CA, USA).

*Processing of raw reads*

Raw reads were processed using a modified STACKS v2 pipeline for RAD-Seq data using a reference genome (Rochette et al., 2019). Reads were quality-checked using *fastqc* (Andrews, 2010). For each sequencing lane and library, paired reads were demultiplexed (assembled by individual), filtered for low quality and missing bases (-c -q) and trimmed to remove barcodes using *process-radtags* in STACKS. For each individual, PCR duplicates were removed using *clone_filter* in STACKS. The remaining sequence data was then merged within individuals across sequencing lanes. Sequences within individuals were aligned to a *C. americana* reference genome (haploid chromosome-level assembly based on the Appalachian clade population VA73 using PacBio HiFi reads; Lopez-Caamal et al., under review at GBE). Based on the alignment, RAD loci across the whole sampling were assembled in a catalogue using *gstacks* in STACKS, resulting in 1,566,971 RAD loci comprising 924,427,374 forward reads and 748,416,814 matched paired-end reads, for an average coverage of 31.2x per individual (SD = 9.7x; range: 8.8x – 80.6x). RAD loci averaged 465bp in length with a mean insert length of 72bp, for a total of 615 Mbp sequenced, covering about ~36% of the 1.7 Gbp haploid reference genome. This catalogue was used to generate the single nuclear polymorphism (SNP) sets for the selected individuals.

*Genotyping for population structure and phylogenetic analyses*

We called genotypes for the 184 individuals as variant call format (VCF) using *populations* in STACKS, retaining SNPs that were present for at least half of the individuals within each population (-r 0.5, not taking into account populations where the site was entirely missing) and for 90% of individuals overall (-R 0.9), with a minor allele count of at least 5 across the 184 samples (--min-mac 5, equivalent to a minor allele frequency of ca. 0.01). The resulting 59,044 SNPs were filtered further to exclude 2,837 SNPs with an average read depth of less than 20 or greater than 70 per individual (average coverage + 4*√average coverage, following (Li, 2014) (average before filtering = 43.4) using VCFTools (Danecek et al., 2011), as well as to exclude 3,751 variants resulting from indels using BCFTools (Danecek et al., 2021). SNPs were further pruned for LD using plink2 (Chang et al., 2015) with a window size of 50kpb and a correlation coefficient of 0.5. This resulted in a final set of 40,195 high quality SNPs sampled from a total sequence length of 5,077,766 bp, with an average of one SNP per ~126 bp. For the population structure and estimated effective migration surfaces (EEMS) analyses, the five samples from the Appalachian clade outgroup population (VA73) were excluded. For the EEMS analysis, we further excluded 177 SNPs that were heterozygous in all individuals and identified as monomorphic by the analysis, violating assumptions.

*Phylogenetic analysis*

We constructed a phylogenetic tree based on the SNP set including the outgroup (Appalachian clade population VA73) using TreeMix v1.12 (Pickrell & Pritchard, 2012), to account for the signatures of admixture identified by inferring population structure (see below). We ran TreeMix with 0 to 10 migration edges (m, i.e. 2x the number of genetic clusters found by sNMF, see Results), 10 iterations per *m* each using a random seed, and a block size (k) of 100. The optimal number of migration edges was inferred using the package *OptM* in R (Fitak, 2021) with the *evanno* method (Evanno et al., 2005), as the *m* with highest second-order rate of change ($\Delta m$, Method S1.2 Table A). For the best *m*, we ran 100 parallel bootstraps. We then generated a consensus tree and assigned bootstraps following the pipeline detailed for the *BITE* package in R (Milanesi et al., 2017).

**Method S2 Table A: Second-order rate of change in likelihood ($\Delta m$) for each migration edge (m)**

| m | $\Delta m$ |
|---|---|
| 0 | NA |
| **1** | **5.228** |
| 2 | 0.296 |
| 3 | 0.625 |
| 4 | 0.303 |
| 5 | 0.110 |
| 6 | 0.117 |
| 7 | 0.472 |
| 8 | 0.279 |
| 9 | 0.504 |
| 10 | NA |

Value in bold represent the best m (highest $\Delta m$)

*Population structure analysis*

We inferred population structure by determining individual-level genotype clustering following an admixture analysis using sparse nonnegative matrix factorization (sNMF) implemented in the *LEA* package in *R* (Frichot & François, 2015). The *snmf* function was run assuming 1 to 36 ancestral populations (*K*), 10 independent runs per *K* (repetitions), and a maximum of 10,000 iterations. The best K was chosen as the one with lowest cross-entropy across runs (Fig. S2.1A, Method S1.2 Table B). For the best *K*, we identified the best run as the one with lowest cross-entropy (Method S1.2 Table B). For the best run of the best K, population-average admixture coefficients were calculated for each genetic cluster from the admixture coefficients of individuals in the *Q*-matrix (function *Q* in LEA) produced by *snmf* (Table S2.2). A genetic cluster was assigned to each population as the cluster with largest average admixture coefficient (Table S2.2).

**Method S2 Table B: Cross-entropy per ancestral populations *K* and independent analysis runs for each *K* of the sNMF population structure analysis**

| Run | *K*=1 | *K*=2 | *K*=3 | *K*=4 | *K*=5 | *K*=6 | *K*=7 | *K*=8 | *K*=9 | *K*=10 | *K*=11 | *K*=12 |
|---|---|---|---|---|---|---|---|---|---|---|---|---|
| 1 | 0.2926 | 0.2819 | 0.2794 | 0.2772 | 0.2782 | 0.2772 | 0.2781 | 0.2789 | 0.2798 | 0.2802 | 0.2822 | 0.2839 |
| 2 | 0.2921 | 0.2814 | 0.2788 | 0.2767 | 0.2780 | 0.2777 | 0.2782 | 0.2785 | **0.2785** | **0.2791** | 0.2809 | 0.2815 |
| 3 | **0.2909** | **0.2802** | 0.2782 | **0.2760** | **0.2756** | 0.2766 | 0.2774 | 0.2786 | 0.2799 | 0.2794 | 0.2813 | 0.2829 |
| 4 | 0.2927 | 0.2821 | 0.2798 | 0.2780 | 0.2772 | 0.2787 | 0.2785 | 0.2801 | 0.2807 | 0.2810 | 0.2839 | 0.2849 |
| 5 | 0.2916 | 0.2808 | 0.2784 | 0.2762 | 0.2771 | 0.2793 | 0.2771 | 0.2784 | 0.2786 | 0.2804 | 0.2802 | **0.2805** |
| 6 | 0.2915 | 0.2806 | 0.2784 | 0.2761 | 0.2759 | 0.2767 | 0.2768 | **0.2775** | 0.2792 | 0.2792 | 0.2817 | 0.2823 |
| 7 | 0.2918 | 0.2810 | 0.2792 | 0.2786 | 0.2775 | 0.2779 | 0.2785 | 0.2797 | 0.2792 | 0.2801 | 0.2808 | 0.2819 |
| 8 | 0.2916 | 0.2809 | **0.2782** | 0.2762 | 0.2758 | **0.2763** | **0.2766** | 0.2787 | 0.2785 | 0.2799 | 0.2817 | 0.2846 |
| 9 | 0.2917 | 0.2811 | 0.2785 | 0.2763 | 0.2759 | 0.2771 | 0.2776 | 0.2792 | 0.2797 | 0.2800 | 0.2808 | 0.2818 |
| 10 | 0.2917 | 0.2810 | 0.2784 | 0.2763 | 0.2763 | 0.2771 | 0.2778 | 0.2779 | 0.2795 | 0.2804 | **0.2796** | 0.2827 |

| Run | *K*=13 | *K*=14 | *K*=15 | *K*=16 | *K*=17 | *K*=18 | *K*=19 | *K*=20 | *K*=21 | *K*=22 | *K*=23 | *K*=24 |
|---|---|---|---|---|---|---|---|---|---|---|---|---|
| 1 | 0.2855 | 0.2869 | 0.2888 | 0.2896 | 0.2914 | 0.2938 | 0.2935 | 0.2957 | 0.2976 | 0.3008 | 0.3032 | 0.3051 |
| 2 | 0.2841 | 0.2863 | 0.2865 | 0.2910 | 0.2884 | 0.2911 | 0.2942 | 0.2962 | 0.2981 | 0.3016 | 0.3016 | 0.3045 |
| 3 | 0.2838 | **0.2832** | 0.2858 | 0.2876 | 0.2900 | 0.2914 | **0.2917** | 0.2958 | 0.2965 | 0.3002 | 0.3003 | 0.3020 |
| 4 | 0.2865 | 0.2845 | 0.2877 | 0.2887 | 0.2893 | 0.2903 | 0.2946 | 0.2974 | 0.2996 | 0.3013 | 0.3059 | 0.3049 |
| 5 | 0.2832 | 0.2855 | 0.2861 | **0.2863** | 0.2889 | 0.2913 | 0.2928 | **0.2937** | 0.2971 | 0.3006 | 0.3031 | 0.3046 |
| 6 | **0.2824** | 0.2844 | 0.2877 | 0.2888 | 0.2898 | **0.2894** | 0.2941 | 0.2952 | 0.2980 | **0.2988** | **0.3002** | 0.3030 |
| 7 | 0.2834 | 0.2851 | 0.2884 | 0.2895 | 0.2904 | 0.2910 | 0.2942 | 0.2949 | 0.2984 | 0.2997 | 0.3031 | 0.3055 |
| 8 | 0.2838 | 0.2850 | 0.2872 | 0.2873 | **0.2875** | 0.2922 | 0.2937 | 0.2965 | 0.2981 | 0.2994 | 0.3022 | **0.3017** |
| 9 | 0.2833 | 0.2874 | **0.2849** | 0.2871 | 0.2892 | 0.2914 | 0.2921 | 0.2963 | **0.2957** | 0.3011 | 0.3015 | 0.3020 |
| 10 | 0.2845 | 0.2853 | 0.2858 | 0.2889 | 0.2901 | 0.2923 | 0.2930 | 0.2950 | 0.2981 | 0.3005 | 0.3035 | 0.3023 |

| Run | *K*=25 | *K*=26 | *K*=27 | *K*=28 | *K*=29 | *K*=30 | *K*=31 | *K*=32 | *K*=33 | *K*=34 | *K*=35 | *K*=36 |
|---|---|---|---|---|---|---|---|---|---|---|---|---|
| 1 | 0.3063 | 0.3068 | 0.3114 | 0.3118 | 0.3181 | 0.3161 | 0.3149 | 0.3219 | 0.3244 | 0.3252 | 0.3293 | 0.3357 |
| 2 | 0.3056 | 0.3108 | 0.3105 | 0.3132 | 0.3163 | 0.3165 | **0.3182** | 0.3238 | 0.3255 | 0.3275 | 0.3317 | 0.3349 |
| 3 | 0.3085 | 0.3097 | 0.3087 | 0.3146 | **0.3120** | 0.3155 | 0.3186 | **0.3201** | 0.3225 | **0.3236** | 0.3316 | 0.3281 |
| 4 | 0.3065 | 0.3106 | 0.3123 | 0.3168 | 0.3197 | 0.3202 | 0.3216 | 0.3239 | 0.3273 | 0.3304 | 0.3342 | 0.3353 |
| 5 | 0.3080 | 0.3075 | 0.3088 | 0.3139 | 0.3143 | 0.3174 | 0.3186 | 0.3232 | 0.3229 | 0.3261 | 0.3291 | 0.3298 |
| 6 | 0.3055 | 0.3075 | **0.3073** | 0.3109 | 0.3137 | 0.3157 | 0.3189 | 0.3220 | 0.3260 | 0.3265 | **0.3258** | 0.3360 |
| 7 | 0.3068 | 0.3114 | 0.3087 | 0.3127 | 0.3155 | 0.3212 | 0.3185 | 0.3211 | 0.3251 | 0.3261 | 0.3295 | 0.3357 |
| 8 | 0.3090 | 0.3090 | 0.3088 | **0.3108** | 0.3148 | 0.3220 | 0.3178 | 0.3226 | 0.3239 | 0.3252 | 0.3294 | **0.3267** |
| 9 | 0.3063 | **0.3064** | 0.3079 | 0.3133 | 0.3129 | **0.3151** | 0.3150 | 0.3261 | **0.3222** | 0.3299 | 0.3272 | 0.3332 |
| 10 | **0.3038** | 0.3069 | 0.3109 | 0.3120 | 0.3135 | 0.3173 | 0.3188 | 0.3229 | 0.3271 | 0.3268 | 0.3292 | 0.3284 |

For each K, the best run – with the lowest cross-entropy – is indicated in bold. The best *K* overall is indicated in gray shading.

*Estimating Effective Migration Surfaces (EEMS) analysis*

To estimate levels of migration, we performed the EEMS analysis (Petkova et al., 2016) using the python package *feems* (Marcus et al., 2021). We conducted the analysis following the recommended workflow detailed in the documentation (https://github.com/Novembrelab/feems, accessed 18/08/2025), except that we used a custom grid generated with the *dggridR* package in R (Fuller Aperture 4 Triangular Grid, resolution 10; (Barnes & Sahr, 2017) with a higher resolution than the default grid provided with *feems*. We performed a leave-one-out cross-validation for the smoothing regularization parameter λ by running *feems* on 20 values of λ ranging from $1e^{-6}$ to $1e^{2}$. This step identifies the model parametrization with the lowest cross-validation error, as values of λ too high or too low can lead to under or overfitting of the model (Marcus et al., 2021). We chose a λ of ~ 0.3 for the final model (Method S1.2 Table C), as lower λ lead to only marginally reduced error and produced similar results (not shown).

**Method S2 Table C: Cross-validation error for each value of the smoothing regularization parameter λ**

| Cross-validation error | λ |
|---|---|
| 0.05527 | 100.000000 |
| 0.05534 | 37.926902 |
| 0.05535 | 14.384499 |
| 0.05518 | 5.455595 |
| 0.05503 | 2.069138 |
| 0.05496 | 0.784760 |
| **0.05490** | **0.297635** |
| 0.05492 | 0.112884 |
| 0.05489 | 0.042813 |
| 0.05490 | 0.016238 |
| 0.05489 | 0.006158 |
| 0.05489 | 0.002336 |
| 0.05489 | 0.000886 |
| 0.05490 | 0.000336 |
| 0.05490 | 0.000127 |
| 0.05490 | 0.000048 |
| 0.05490 | 0.000018 |
| 0.05490 | 0.000007 |
| 0.05490 | 0.000003 |
| 0.05490 | 0.000001 |

Value in bold represents the λ chosen for analysis (lowest value after which the cross-validation error is negligeable).
