## Appendix 2 for "Heterogeneous evolutionary history defines the rear edge of the North American herb *Campanula americana*"

**Appendix 2: Supplementary Results**

**Supplementary tables**

**Table S2.1: Model comparisons testing the linear and 2nd or 3rd degree quadratic effect of latitude**

| Dependent variable | *N* | *AICc* linear | *AICc* square | *AICc* cubic |
|---|---|---|---|---|
| LGM suitability | 763 | 541.44 | 490.86 | **485.76** |
| Present-day suitability | 763 | 242.18 | **214.15** | 214.08 |
| Present-day connectivity | 763 | 357.23 | 283.95 | **272.22** |

The best model (bold) was identified by the lowest AICc value. For models with similar AICc (|ΔAICc|<2), the simpler model was chosen.

**Table S2.2: Populations used, genetic cluster assignment and population-average admixture per genetic cluster**

| Population | Latitude [°N] | Longitude [°E] | Genetic cluster | Clade | Population-average admixture | | | | |
|---|---|---|---|---|---|---|---|---|---|
| | | | | | Refugia E | Refugia W | Core | Expansion N | Expansion S |
| FL83 | 30.56468 | -84.95982 | Refugia E | Refugia | **0.9414** | 0.0001 | 0.0507 | 0.0001 | 0.0077 |
| FL81 | 30.81165 | -85.22535 | Refugia E | Refugia | **0.9996** | 0.0001 | 0.0001 | 0.0001 | 0.0001 |
| AL2 | 31.54792 | -87.51453 | Refugia E | Refugia | **0.7485** | 0.0925 | 0.1366 | 0.0163 | 0.0061 |
| MS6 | 31.99803 | -89.35603 | Refugia W | Refugia | 0.0119 | **0.8175** | 0.0001 | 0.1267 | 0.0438 |
| AL23 | 32.19395 | -86.78484 | Refugia E | Refugia | **0.9293** | 0.0198 | 0.0376 | 0.0064 | 0.0069 |
| AL8 | 32.25610 | -86.49330 | Refugia E | Refugia | **0.9437** | 0.0074 | 0.0238 | 0.0111 | 0.0140 |
| AL22 | 32.50376 | -87.50489 | Refugia W | Refugia | 0.2619 | **0.4998** | 0.1982 | 0.0089 | 0.0312 |
| AL19 | 32.60699 | -88.19192 | Refugia W | Refugia | 0.0004 | **0.9993** | 0.0001 | 0.0001 | 0.0001 |
| AL79 | 32.92932 | -88.20820 | Refugia W | Refugia | 0.0078 | **0.9126** | 0.0546 | 0.0018 | 0.0231 |
| AL21 | 33.49109 | -86.79829 | Core | Refugia | 0.1562 | 0.0773 | **0.6886** | 0.0189 | 0.0589 |
| MS9 | 33.92778 | -89.01710 | Refugia W | Refugia | 0.0001 | **0.9575** | 0.0016 | 0.0329 | 0.0079 |
| OK61 | 33.94639 | -94.56694 | Expansion S | Expansion | 0.0003 | 0.0001 | 0.0001 | 0.0001 | **0.9994** |
| MS8 | 34.40427 | -88.83093 | Refugia W | Refugia | 0.0107 | **0.7522** | 0.0820 | 0.1303 | 0.0248 |
| AL7 | 34.47732 | -87.53973 | Core | Refugia | 0.0693 | 0.1004 | **0.8018** | 0.0033 | 0.0252 |
| GA1 | 34.60074 | -84.69664 | Core | Refugia | 0.1363 | 0.0001 | **0.8434** | 0.0001 | 0.0201 |
| ALBG | 34.65346 | -86.51638 | Core | Refugia | 0.0078 | 0.0041 | **0.9879** | 0.0001 | 0.0001 |
| TN3 | 35.30932 | -90.06770 | Expansion N | Expansion | 0.0373 | 0.1468 | 0.1683 | **0.4620** | 0.1856 |
| TN2 | 35.74976 | -85.38068 | Core | Refugia | 0.0949 | 0.0061 | **0.8851** | 0.0001 | 0.0138 |
| OK1 | 36.08091 | -95.05494 | Expansion S | Expansion | 0.0537 | 0.0196 | 0.0841 | 0.0001 | **0.8425** |
| TN34 | 36.08222 | -86.29611 | Core | Expansion | 0.0218 | 0.0280 | **0.9078** | 0.0247 | 0.0177 |
| AR2 | 36.15235 | -94.30464 | Expansion S | Expansion | 0.0001 | 0.0010 | 0.0001 | 0.0023 | **0.9965** |
| KY5 | 37.36005 | -84.77186 | Core | Expansion | 0.0233 | 0.0396 | **0.8790** | 0.0415 | 0.0165 |
| KY1 | 38.09215 | -84.98688 | Core | Expansion | 0.0291 | 0.0598 | **0.8435** | 0.0351 | 0.0325 |
| MO2 | 38.83025 | -92.28509 | Expansion N | Expansion | 0.0218 | 0.0349 | 0.0107 | **0.7583** | 0.1743 |
| KS60 | 39.04742 | -95.68152 | Expansion N | Expansion | 0.0200 | 0.0214 | 0.0120 | **0.5898** | 0.3568 |
| IN5 | 39.14583 | -86.54833 | Core | Expansion | 0.0001 | 0.0132 | **0.6292** | 0.3380 | 0.0195 |
| IN7 | 39.87011 | -86.16020 | Core | Expansion | 0.0001 | 0.0242 | **0.4954** | 0.4663 | 0.0140 |
| OH119 | 39.88500 | -83.99700 | Core | Expansion | 0.0001 | 0.0114 | **0.8581** | 0.1200 | 0.0105 |
| IL10 | 40.61750 | -89.01778 | Expansion N | Expansion | 0.0233 | 0.0313 | 0.0429 | **0.8794** | 0.0231 |
| IN2 | 41.01731 | -85.23872 | Expansion N | Expansion | 0.0001 | 0.0009 | 0.2388 | **0.7601** | 0.0001 |
| IA12 | 41.69417 | -93.67417 | Expansion N | Expansion | 0.0001 | 0.0001 | 0.0001 | **0.9142** | 0.0855 |
| IA17 | 42.46194 | -90.63611 | Expansion N | Expansion | 0.0092 | 0.0110 | 0.0049 | **0.9707** | 0.0043 |
| MI2 | 42.62369 | -85.43706 | Expansion N | Expansion | 0.0001 | 0.0093 | 0.3707 | **0.6198** | 0.0001 |
| WI4 | 43.40972 | -89.63667 | Expansion N | Expansion | 0.0247 | 0.0017 | 0.0094 | **0.9627** | 0.0015 |
| MN8 | 44.02861 | -92.43222 | Expansion N | Expansion | 0.0093 | 0.0049 | 0.0001 | **0.9270** | 0.0587 |
| MN117 | 44.90100 | -93.19200 | Expansion N | Expansion | 0.0076 | 0.0084 | 0.0103 | **0.8603** | 0.1133 |

Genetic clusters (for *K*=5) were assigned for each population as the cluster with highest population-average admixture coefficients (bold).

### Supplementary Figures

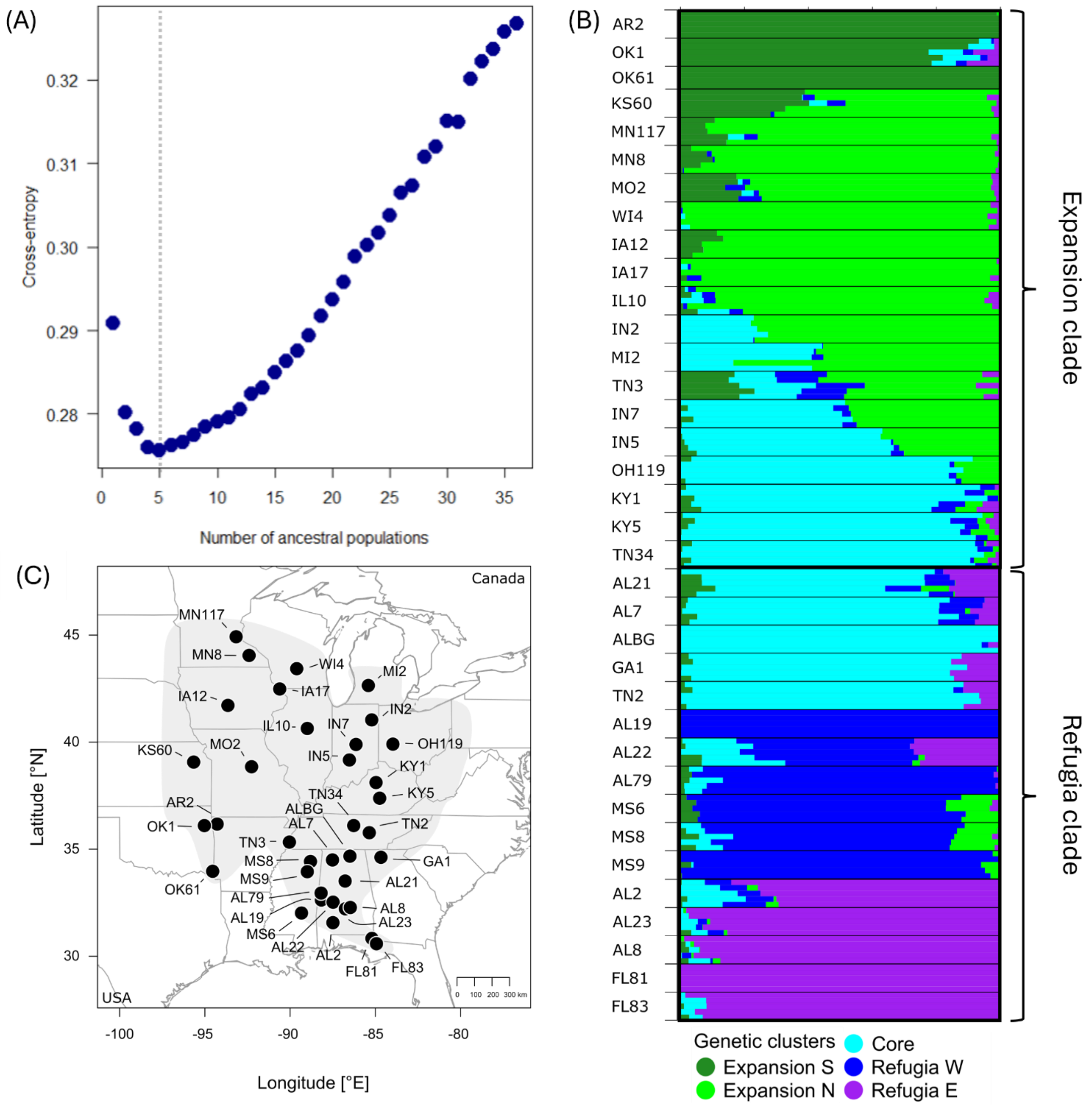


**Figure S2.1: Parametrization of the population structure analysis, individual cluster assignments and populations sequenced**

**(A)** Cross-entropy for each number of ancestral populations $K$, with best $K$ indicated by the vertical dotted line. **(B)** Genetic cluster assignment for each individual for $K = 5$, colors indicate distinct genetic clusters. Samples are sorted by populations, and populations by main genetic cluster (cluster with highest population average admixture). Brackets detail which populations represent the expanded and the refugia clade in the phylogenetic analysis (see Fig. 2). **(C)** Location of the sequenced populations (circles) with population labels, and the distribution of the Western clade (gray shading).
